# Graded Hair Cell Ablation Reveals Functional Redundancy in the Mature Mouse Vestibular System

**DOI:** 10.1101/2025.11.27.691044

**Authors:** Tian Wang, Davood K. Hosseini, Ahmad Mahhoudi, Zahra N. Sayyid, Jun He, Caroline Sit, Zelma G. Iriarte Oporto, Hong Zhu, Wu Zhou, Alan G. Cheng

## Abstract

Inner ear hair cells are mechanoreceptors critical for hearing and balance. Damage to vestibular hair cells causes balance impairment, yet it is unclear whether hair cell loss correlates with vestibular dysfunction. In this study, we ablated hair cells in *Pou4f3^DTR/+^* mice with diphtheria toxin (DT), and found a dose-dependent decrease in hair cell survival in both the macula and crista ampullaris, including loss of type I and II hair cells in the striolar/central and extrastriolar/peripheral regions. Responses to linear acceleration, measured by the translational vestibulo-ocular reflex (tVOR) and vestibular sensory evoked potential (VsEP), were intact with 25% or more hair cell survival in the macula, and diminished only when hair cell survival decreased further. By contrast, rotational vestibulo-ocular reflex (rVOR) responses were significantly reduced with ∼31% hair cell survival in the cristae. Further, single-unit recordings of vestibular afferents from cristae and maculae showed more irregular and reduced firing rates, but only those corresponding to cristae displayed reduced sensitivity to head rotation. Limited hair cell regeneration was observed in the extrastriolar/peripheral regions of both maculae and cristae 6 months post ablation, although no significant recovery of the VORs was observed. Thus, the adult mouse vestibular end organs display different degrees of redundancy, demonstrating robust responses to head rotation and linear acceleration despite loss of most hair cells.

## Introduction

The vestibular system is essential for maintaining balance, spatial orientation, and gaze stabilization. It comprises five pairs of sensory organs in the inner ear: crista ampullaris in the three semicircular canals, which function to detect head rotational acceleration; the two macular/otolith organs, the utricle and saccule, detect linear acceleration and head tilt with respect to gravity. Dysfunction of the vestibular system leads to dizziness, vertigo, and falls, with approximately 35% of US adults aged 40 and older having experienced these symptoms (Agrawal, Carey et al. 2009, Agrawal, Ward et al. 2013, Agrawal, Merfeld et al. 2020).

Inner ear hair cells are mechanosensors essential for vestibular function, and their loss due to factors including aging, ototoxic drugs (e.g., aminoglycosides), and genetic mutations can lead to significant balance disorders (Tsuji, Velazquez-Villasenor et al. 2000, Rauch, Velazquez-Villasenor et al. 2001). Two types of mechanosensory hair cells populate the vestibular organs: type I and type II hair cells, which can be distinguished by molecular markers, cell morphology, patterns of neuronal innervation, and electrophysiological properties (Goldberg 2000, Eatock and Songer 2011, Burns and Stone 2017). To study hair cell degeneration and regeneration of the vestibular system, several ototoxins have been used to induce hair cell loss. A single systemic dose of aminoglycosides can induce hair cell loss in the vestibular organs in guinea pigs, including the cristae ampullaris and macular organs, despite limited efficacy in mice (Lang and Liu 1997, Nakagawa, Yamane et al. 1998). Alternatively, a nitrile toxin such as IDPN (3,3’-iminodipropionitrile) effectively ablates vestibular hair cells but also causes central nervous system toxicity and hyperactive behavior (Llorens, Dememes et al. 1994). The Pou4f3-DTR mouse line is a well-established genetic model for selective hair cell ablation in the inner ear (Lin, Golub et al. 2011, Golub, Tong et al. 2012). In these mice, expression of the human diphtheria toxin receptor (DTR) is driven by the hair cell promoter Pou4f3. Systemic administration of diphtheria toxin (DT) induces rapid and targeted ablation of vestibular hair cells, leading to vestibular dysfunction manifested by abnormalities in behavior and vestibular physiology measurements, including vestibular-evoked potentials (VsEP), vestibulo-ocular reflex (VOR), and single-unit recording of vestibular afferents (Golub, Tong et al. 2012, Jauregui, Scheinman et al. 2024, Lahlou, Zhu et al. 2024).

Vestibular function is classically assessed using a battery of behavioral and physiological assays, each with its own specificity and sensitivity for individual vestibular end-organs. Open field/swimming/rotarod tests assess gross motor function related to balance control but are neither specific nor sensitive for detecting vestibular deficits (Goldberg, Wilson et al. 2012). The rotational vestibulo-ocular reflex (rVOR) measures crista/canal function. In contrast, translational vestibulo-ocular reflex (tVOR) measures macular/otolith function in response to a range of frequencies, the vestibular-sensory evoked potential (VsEP) assesses otolith function, presumably relying on the synchronized activity of calyx afferents that innervate striolar type I hair cells (Jones, Jones et al. 2011, Ono, Keller et al. 2020). In vivo single-unit recordings of vestibular afferents quantitatively assess spontaneous discharge rates, regularity, and dynamic responses to head rotation and translation. Because it is highly specific and sensitive to vestibular deficits, it is considered the “gold standard” for assessing peripheral vestibular function in animal models (Goldberg, Wilson et al. 2012).

Previous studies have demonstrated that near-complete ablation of vestibular hair cells (>90%) in Pou4f3-DTR mice abolishes both rVOR and tVOR responses (Jauregui, Scheinman et al. 2024, Lahlou, Zhu et al. 2024). However, our understanding of how different degrees of damage affect vestibular function is rather limited. In this study, we assessed the relationship between escalating degrees of hair cell loss and vestibular function by administering different doses of DT into adult Pou4f3-DTR mice. In the utricle, we found that VsEP thresholds were maintained at 25% hair cell survival after DT administration, and only decreased after ∼92% hair cell loss. Similarly, tVOR responses remained largely unchanged when only 25% utricular hair cells remained. However, deficits in rVOR responses were detected with 31% hair cell survival in the cristae, and worsened after ∼89% hair cell loss. Finally, single unit recording detected decreased spontaneous firing rates and regularity in both the crista and utricle at 25-31% hair cell survival. Together, these findings suggest that adult mouse vestibular hair cells are redundant, with a higher degree of redundancy in the otolith than crista organs.

## Result

### Strain-dependent animal survival after DT injection

The Pou4f3-DTR mice have been established as a reliable model system to study hair cell ablation in both the cochlear and vestibular organs (Lin, Golub et al. 2011, Golub, Tong et al. 2012). The hair cell promoter Pou4f3 drives expression of the human DTR, allowing selective ablation of hair cells upon DT administration, while preserving supporting cells and innervating neurons. Previous studies using Pou4f3-DTR mice on C57BL/6J, CBA/J, or mixed backgrounds have reported low mortality (<5%) (Golub, Tong et al. 2012, Tong, Strong et al. 2015). We first compared Pou4f3-DTR mice from two different genetic backgrounds by administering DT at 4 weeks of age (DT, one or two doses spaced 48 hours apart, on days 1 and 3). Pou4f3-DTR mice on a FVB background were generated by backcrossing C57BL/6J Pou4f3-DTR mice with wild-type FVB mice for over ten generations. After DT administration, Pou4f3-DTR mice on the FVB background experienced high mortality (>60%). Specifically, survival rates were 38.5% after a single dose of with DT at 15 ng/g (n=13), and further decreased at higher doses (33.0% for 15 ng/g x2 (n=3), 0% with 20 ng/g x2 (n=3), 25 ng/g x2 (n=5) and 35 ng/g x2 (n=5)) (Table 1). Mortality typically occurred within 1-2 days of DT injection, with notable weight loss and dehydration. Subcutaneous administration of lactated Ringer’s solution (3 times daily) and high-calorie gel for two weeks markedly improved animal survival (100% at 15 ng/g (n=8), 81% at 25 ng/g x2 (n=27), 63% at 35 ng/g x2 (n=33), and 50% at 50 ng/g x2 (n=10)) (Table 1). Remarkably, no mortality was observed in DT-injected, supplement-free Pou4f3-DTR mice on a C57BL/6J background (25 ng/g x2 (n=13) and 50 ng/g x2 (n=9)) (Table 1). These findings indicate strain-dependent differences in DT-induced mortality. As higher doses of DT are needed to induce hair cell ablation in the vestibular system than in the cochlea, Pou4f3-DTR mice on a C57BL/6J background may be more suitable for studying vestibular hair cell degeneration/regeneration in adult mice.

### DT induces dose-dependent hair cell loss in the maculae and cristae

To assess the relationship between DT dose/regimen and vestibular hair cell loss, utricles and anterior cristae ampullaris were collected from Pou4f3-DTR mice two weeks after DT injection (Figure 1A and 2A). The area containing Myosin7a+ hair cells showed substantial reduction, with 92.5±6.8%, 87.0±7.3%, 63.6±11.1%, and 83.4±12.0% of wild type controls at DT doses of 15 ng/g x1, 25, 35, and 50 ng/g x2, respectively, compared to age-matched controls (Figure 1B-F, Supplementary figure 1A). In the utricle, hair cell survival in the extrastriola decreased to 58.4±7.4% after low-dose DT treatment (15 ng/g x1) compared to controls. At higher DT doses, hair cell survival further decreased to 26.8±6.8%, 9.5±2.8%, and 19.2±5.2% in a dose-dependent manner (25 ng/g x2, 35 ng/g x2, and 50 ng/g x2) (Figure 1B-G, Table 2). Similarly, hair cell survival in the striola declined more after higher doses of DT treatment (53.7±8.1%, 23.6±8.3%, 6.6±2.9% and 17.1±9.8% at 15 ng/g, 25, 35, and 50 ng/g x2, compared to age-matched controls, respectively (Figure 1B-F, H, Table 2). Similar degree of hair cell loss was observed in organs from the two strains of mice.

**Figure 1.**
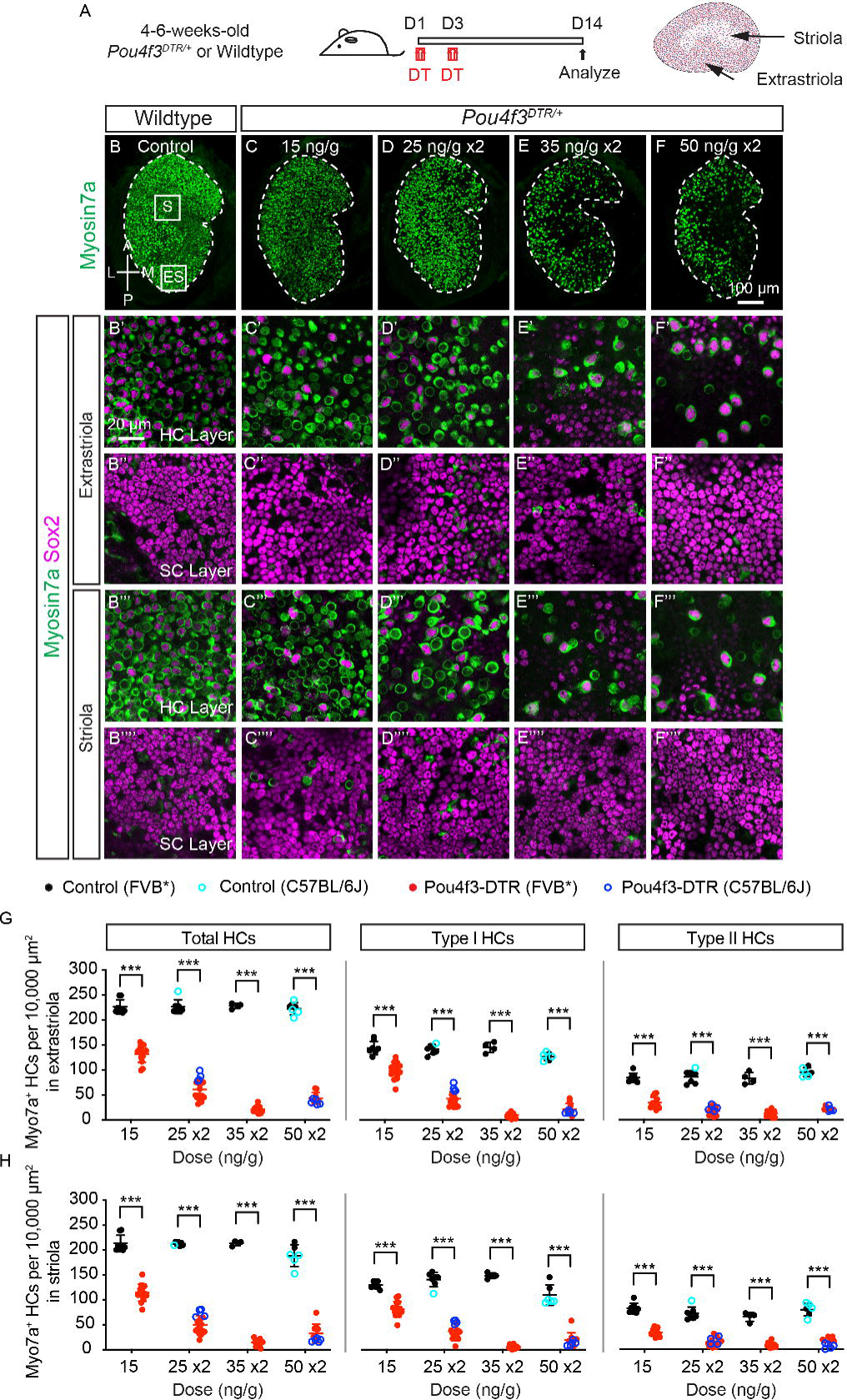
Diphtheria toxin induces dose-dependent hair cell loss in the utricle of adult *Pou4f3^DTR/+^* mice. **A.** Schematic showing diphtheria toxin treatment of 4-6-week-old *Pou4f3^DTR/+^* and wild-type mice on Day 1 and Day 3, and inner ear tissues were analyzed 2 weeks later. **B-F.** Low-magnification images of the utricle labeled for Myosin7a. Samples include wild-type and *Pou4f3^DTR/+^*mice treated with various DT doses (15, 25 x2, 35 x2, and 50 x2 ng/g). A small but significant reduction in the sensory epithelium (Myosin7a^+^) was observed with all DT doses tested. **B’-F’’’’.** High-magnification images of the utricle showing a dose-dependent Myosin7a⁺ hair cell loss after DT treatment, while supporting cells remain unaffected in both extrastriola (**B’**-**B’’**) and striola (**B’’’**-**B’’’’**). **G-H.** Quantification of Myosin7a^+^ HCs showing a significant reduction in the density of total HCs and each HC subtype in the extrastriolar (**G**) and striolar (**H**) regions after DT treatment specified doses in both FVB (black and red dots) and C57BL/6J (cyan and blue circles) mice. Greater losses were noted at higher DT doses. Data shown as mean±SD, and were analyzed using two-tailed, unpaired Student *t* tests. ***p<0.001, **p<0.01, *p<0.05. n=8, 7, 4, and 6 mice for wild type, and n=14, 18, 13, and 10 mice for *Pou4f3^DTR/+^*, treated with 15, 25 x2, 35 x2, and 50 x2 ng/g DT, respectively. Scale bars: 100 μm in (B-F), 20 μm in (B’-F’’’’).

**Figure 2.**
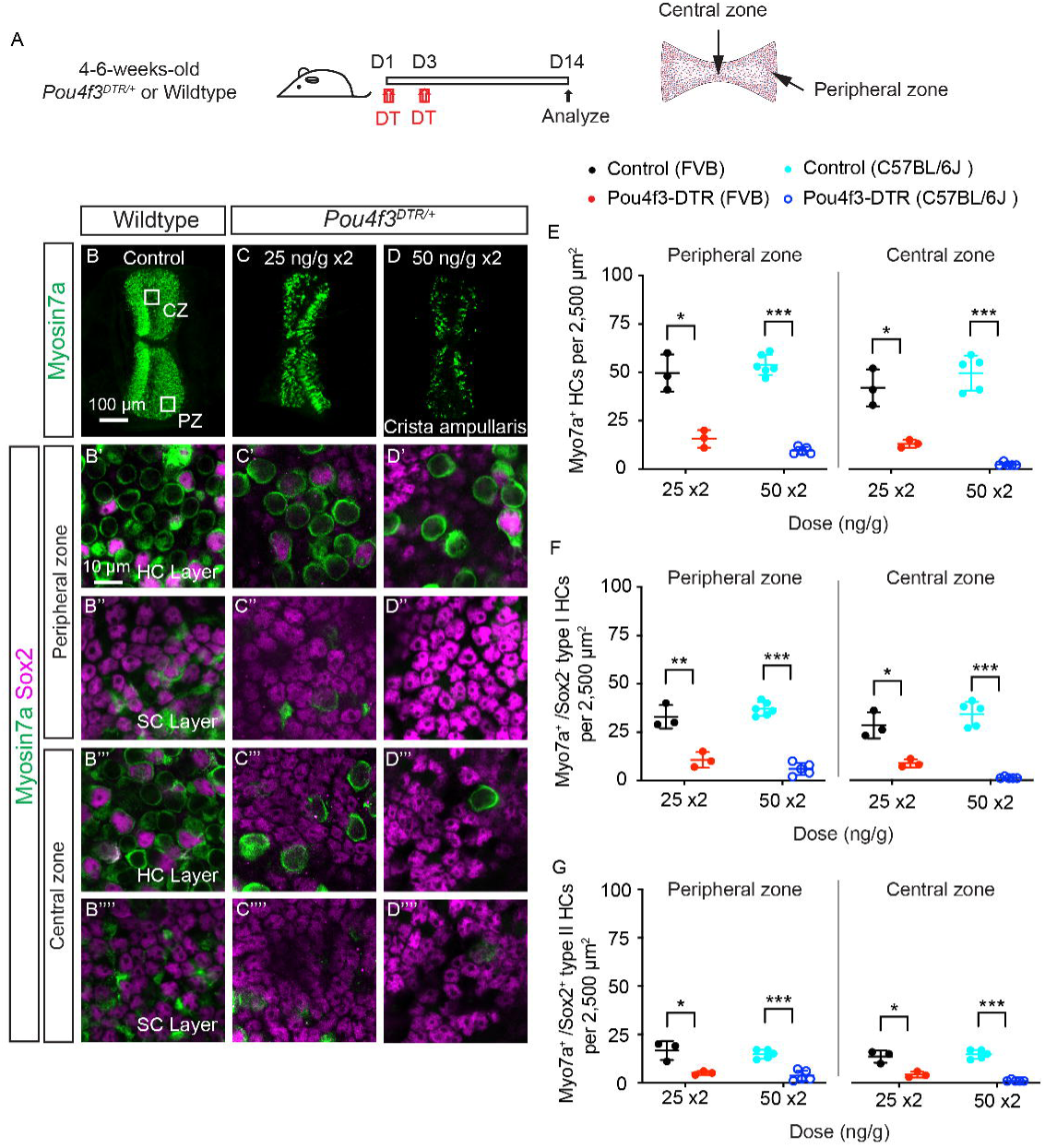
Diphtheria toxin induces dose-dependent hair cell loss in the crista ampullaris of adult *Pou4f3^DTR/+^* mice. **A.** Schematic showing DT administration to 4-6-week-old *Pou4f3^DTR/+^* and wild-type mice, and inner ear tissues were harvested 2 weeks later. **B-D.** Low-magnification images of the crista ampullaris showing marked reduction of Myosin7a^+^ hair cells after DT treatment of *Pou4f3^DTR/+^*mice (25 x2 and 50 x2 ng/g). **B’-D’’’’.** High-magnification images of the crista ampullaris showing dose-dependent Myosin7a⁺ hair cell loss in both the peripheral (**B’**-**B’’**) and central zones (**B’’’**-**B’’’’**),, while supporting cells appeared intact. **E-G.** Quantitative analysis shows a significant reduction of total Myosin7a^+^ HCs (**E**), type I HCs (**F**), and type II HCs (**G**) in peripheral (left panels) and central (right panels) zones following DT treatment in both FVB (black and red dots) and C57BL/6J (cyan and blue circles) mice. Data shown as mean±SD, and were analyzed using two-tailed, unpaired Student *t* tests. ***p<0.001, **p<0.01, *p<0.05. n=3 and 6 mice for wild type, and n=3 and 5 mice for *Pou4f3^DTR/+^*, treated with 25 x2 and 50 x2 ng/g DT, respectively. Scale bars: 100 μm in (B-D); 10 μm in (B’-D’’’’).

Both type I and II hair cells reside in the striolar and extrastriolar regions of the utricle. Using Myosin7a and Sox2 to differentially mark type I and II hair cells, we found that extrastriolar type I hair cells (Myosin7a+/Sox2-) were markedly reduced by DT treatment (68.1±11.7 %, 30.0±10.3%, 6.3±3.1%, and 16.6±9.2% at 15 ng/g x1, 25, 35, and 50 ng/g x2, compared to controls, respectively). Similarly, DT treatment decreased striolar type I hair cells to 62.7±11.9%, 25.8±10.3%, 3.8±2.2%, and 17.4±13.7% (Figure 1B-H, Table 2). Finally, DT administration led to loss of type II hair cells (Myosin7a+/Sox2+) in the extrastriola (40.5±13.0%, 21.4±9.7%, 15.2±7.2%, and 22.8±4.3%, respectively), and striola (40.5±8.5%, 19.4±11.7%, 12.9±7.4%, and 16.8±11.0%, respectively (Figure 1B-H, Table 2). At all DT doses tested, supporting cell numbers remained unchanged compared to controls (Figure 1B’’-F’’, B’’’’-F’’’’, Supplementary figure 1B-C). These results suggest that DT induces a dose-dependent loss of type I and II hair cells in both the striolar and extrastriolar regions of the adult mouse utricle, with both hair cell subtypes showing similar susceptibility.

In the anterior cristae of DT-injected Pou4f3-DTR mice (25 ng/g x2, both FVB and C57BL/6J background), we found a lower hair cell survival than undamaged controls (31.5±9.1% in the peripheral zone and 31.0±4.8% in the central zone, respectively, Figure 2A-C, E, Table 2). At a higher dose of DT (50 ng/g x2), hair cell survival further decreased (17.8±3.4% and 4.8±1.8% in the peripheral and central zones, respectively (Figure 2B, D, E, Table 2). After DT treatment (25 ng/g x2), the survival of Myosin7a+/Sox2- type I hair cells declined to 32.3±12.2% and 30.6±7.3% in the peripheral and central zones, respectively. At a higher dose (50 ng/g x2), type I hair cell survival declined to 16.2±8.5% and 3.5±1.3% in the peripheral and central zones, respectively (Figure 2B’-D’’’’, F, Table 2). Similarly, type II hair cell survival significantly decreased after DT (25 and 50 ng/g x2) treatment: only 30.0±6.0% and 24.3±18.2% hair cells remained in the peripheral zone, and 31.7±11.1% and 8.1±3.0% in the central zone compared to controls (Figure 2B’-D’’’’, G, Table 2). These data indicate that DT injection in Pou4f3-DTR mice induced a dose-dependent hair cell loss in both the maculae and cristae, with a significant reduction in both type I and type II hair cells across the extrastriolar/striolar and peripheral/central zones.

### Spontaneous hair cell regeneration in the utricle and crista ampullaris

Previous studies have reported spontaneous hair cell regeneration after severe damage to the utricle and cristae in the Pou4f3-DTR mice (Golub, Tong et al. 2012, Gonzalez-Garrido, Pujol et al. 2021, Jauregui, Scheinman et al. 2024). To determine if the regenerative capacity is altered by different degrees of hair cell loss, we administered DT at two doses (25 and 50 ng/g x2) to 4- to 6-week-old Pou4f3-DTR mice (Figure 3A, 4A). In the utricle six months post DT injection, hair cell density in the extrastriola is significantly higher than that at 2 weeks post DT injection (by 51.8±8.8% in the 25 ng/g x2 group, 130.3±7.6 versus 85.9±9.3 per 10,000 µm^2^, p<0.05, and by 117.6±21.0% in the 50 ng/g x2 group, 78.3 ± 7.6 versus 36.0 ± 5.2 per 10,000 µm^2^, p<0.001, Figure 3B-F’’’’, G–H, Table 3). When hair cell subtypes were segregated, we found that only type II hair cells (Myosin7a+/Sox2+) significantly increased (22.5±5.2 to 54.0 ±2.6 per 10,000 µm^2^, p<0.01, and 20.9±4.5 to 57.3±6.7 per 10,000 µm^2^, p<0.001, in the 25 ng/g x2 and 50 ng/g x2 groups, respectively), while type I hair cells (Myosin7a+/Sox2-) showed no significant change (Figure 3G-H, Table 3). Although the striola showed an increasing trend in total and type II hair cell density over time (Figure 3C-H, Table 3), these changes were not statistically significant. Notably, hair cell density in both the extrastriola and striola remained significantly lower than in the undamaged controls, indicating limited spontaneous regeneration, consistent with previous reports (Golub, Tong et al. 2012, Sayyid, Wang et al. 2019). These findings reveal a dose-dependent regeneration of type II hair cells in the extrastriolar region of the adult mouse utricle.

**Figure 3.**
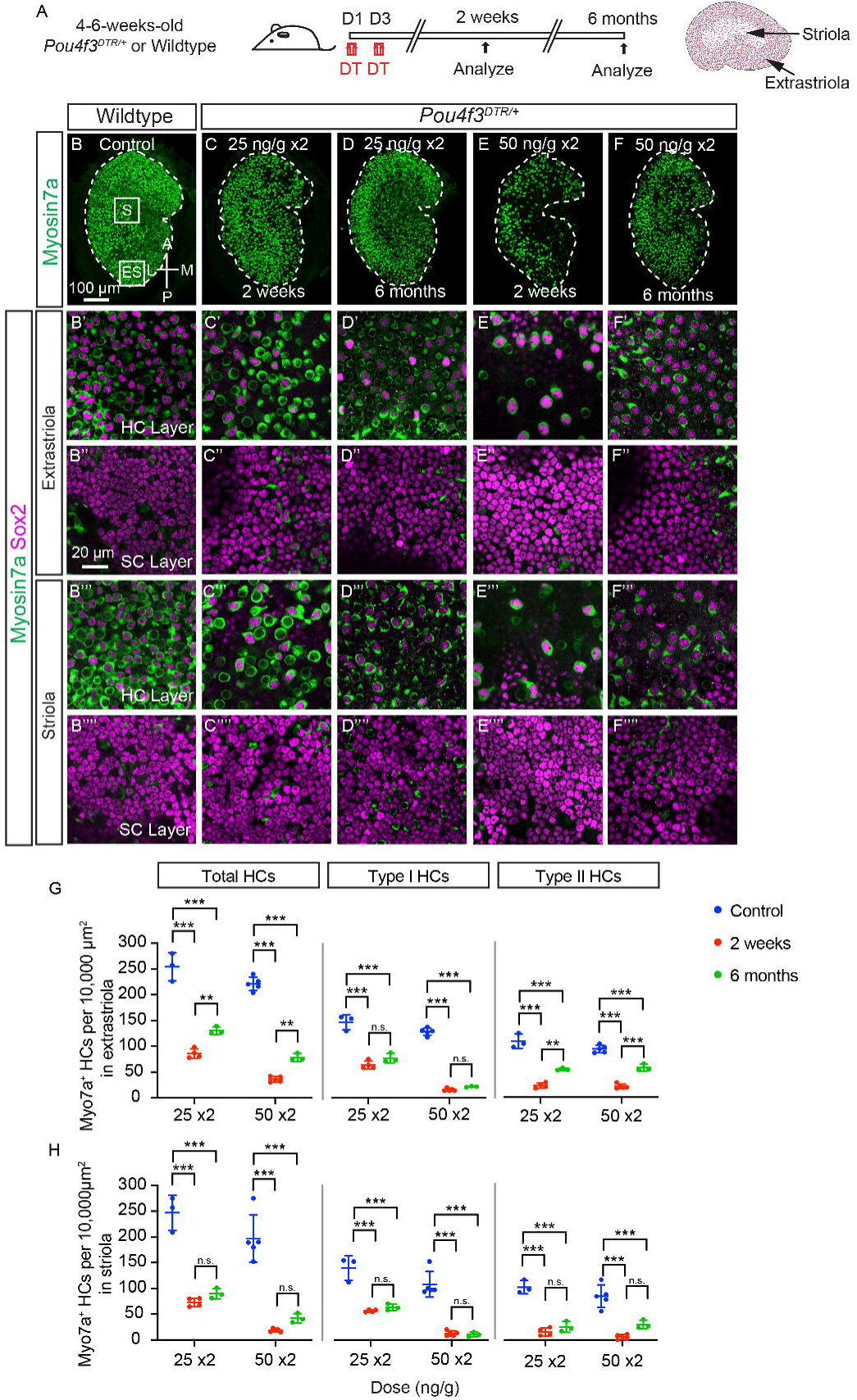
Hair cell regeneration in the utricle following diphtheria toxin damage. **A.** Schematic showing the experimental paradigm of DT treatment of 4-6-week-old *Pou4f3^DTR/+^* transgenic and wild-type mice. **B-F.** Low-magnification images of the utricle showing modest regeneration of Myosin7a^+^ hair cells 6 months after damage by DT treatment at 25 or 50 ng/g x2. **B’-F’’’’.** High-magnification images showing high density of Myosin7a^+^ hair cells at 6 months. **G-H.** Quantification of Myosin7a^+^ hair cell density shows a significant increase in total HC and type II hair cells at 6 months post damage in the extrastriolar region. There was no significant increase in hair cells in the striolar region or type I hair cells in the extrastriolar region. Hair cell numbers in both regions remained significantly lower than in wild-type controls. Data shown as mean±SD, and were analyzed using two-way ANOVA test with ordinary multiple comparison. ***p<0.001, **p<0.01, *p<0.05. n=3 and 5 mice for wild type, n=4 and 5, 3 and 3 mice for *Pou4f3^DTR/+^* at 2 weeks and 6 months, treated with 25 x2 and 50 x2 ng/g DT, respectively. Scale bars: 100 μm in (B–F), 20 μm in (B’–F’’’’).

As described above, DT injection significantly reduced hair cell density throughout the anterior cristae at 2 weeks post-injection. By 6 months post-injection, hair cell density in the peripheral zone significantly increased for the 25 ng/g x2 regimen, but not 50 ng/g x2 (15.7±4.5 to 31.0±5.4 per 2,500 µm^2^, p<0.01, and 9.6±1.8 to 15.0 ± 2.0 per 2,500 µm^2^, respectively, Figure 4B-G, Table 3). Type II hair cells (Myosin7a+/Sox2+) showed a trend of increase in the 25 ng/g x2 group (from 8.7±2.1 to 12.8±6.9 per 2,500 µm^2^), and 50 ng/g x2 group (1.2±0.4 to 2.7±0.6 per 2,500 µm^2^, Table 3). There was no significant change in the density of type I hair cells (Myosin7a+/Sox2-) in either dose (Table 3). Finally, the central zone displayed no significant change in density of total or subtypes of hair cells at both doses (Figure 4C–F′′′′, H, Table 3). Together, these findings suggest that hair cells partially regenerate in the peripheral zone of the adult crista ampullaris after moderate but not severe damage.

**Figure 4.**
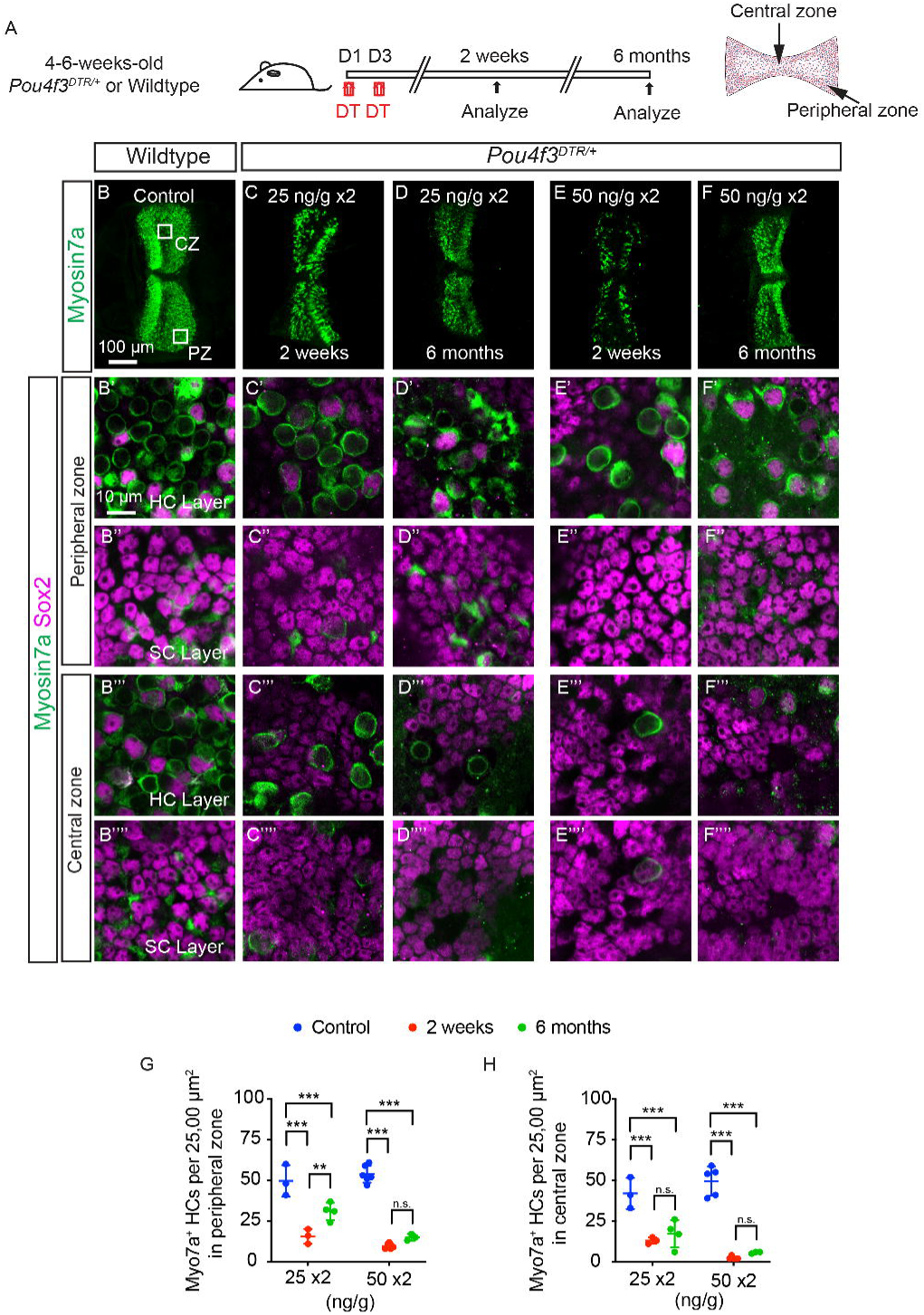
Dose-dependent regenerative response in the crista ampullaris following diphtheria toxin damage. **A.** Schematic showing regimen used for DT treatment of 4-6-week-old *Pou4f3^DTR/+^*and wild-type mice. **B-F.** Low-magnification images showing DT-induced hair cell loss in the crista ampullaris **B’-F’’’’.** High-magnification images showing a higher density of Myosin7a⁺ hair cells in the peripheral zone, but not the central zone, at 6 months post DT treatment. **G-H.** Quantification of Myosin7a^+^ HCs showing significant hair cell regeneration in the peripheral (G) but not central (H) zones following DT treatment at 25 ng/g x2 but not at 50 ng/g x2. Data shown as mean±SD, and were analyzed using one-way ANOVA test. ***p<0.001, **p<0.01, *p<0.05. n=3 and 6 mice for wild type, n=3 and 5, 4 and 3 mice for *Pou4f3^DTR/+^* at 2 weeks and 6 months, treated with 25 x2 and 50 x2 ng/g DT, respectively. Scale bars: 100 μm in (B-F), 10 μm in (B’-F’’’’).

### Vestibular dysfunction after DT-induced hair cell ablation: VsEPs and VORs

VsEP measures otolith function and is particularly dependent on the synchronized activity of calyx afferents that innervate striolar type I hair cells (Jones, Jones et al. 2011, Ono, Keller et al. 2020). After DT administration at 2 weeks, VsEP thresholds, amplitudes, and latencies of the Pou4f3-DTR mice remained unchanged at low dose (15 ng/g x1), with ∼54% hair cell survival in striola (Figure 5A-E). At 25ng/g x2, with ∼24% hair cell survival in striola, VsEP thresholds were not significantly different, although latencies were increased, and amplitudes were reduced in Pou4f3-DTR mice (p<0.001, <0.05, Figure 5B-E). Notably, only 3 out of 13 mice exhibited elevated VsEP thresholds at this dose. In contrast, when only ∼7% of hair cells survived in striola (35 ng/g x2), VsEP thresholds and latencies were significantly increased, and amplitudes were significantly decreased (p<0.001, <0.001, <0.001). At the highest dose (35 ng/g x2), 53.8% (7 out of 13) of the animals showed elevated VsEP thresholds, and one of them had a complete loss of VsEP responses (Figure 5B-E). These data suggest that VsEP responses are more frequently abnormal in mice with lower hair cell survival as a result of higher DT doses, although responses are still detectable with only <10% hair cells remaining.

**Figure 5.**
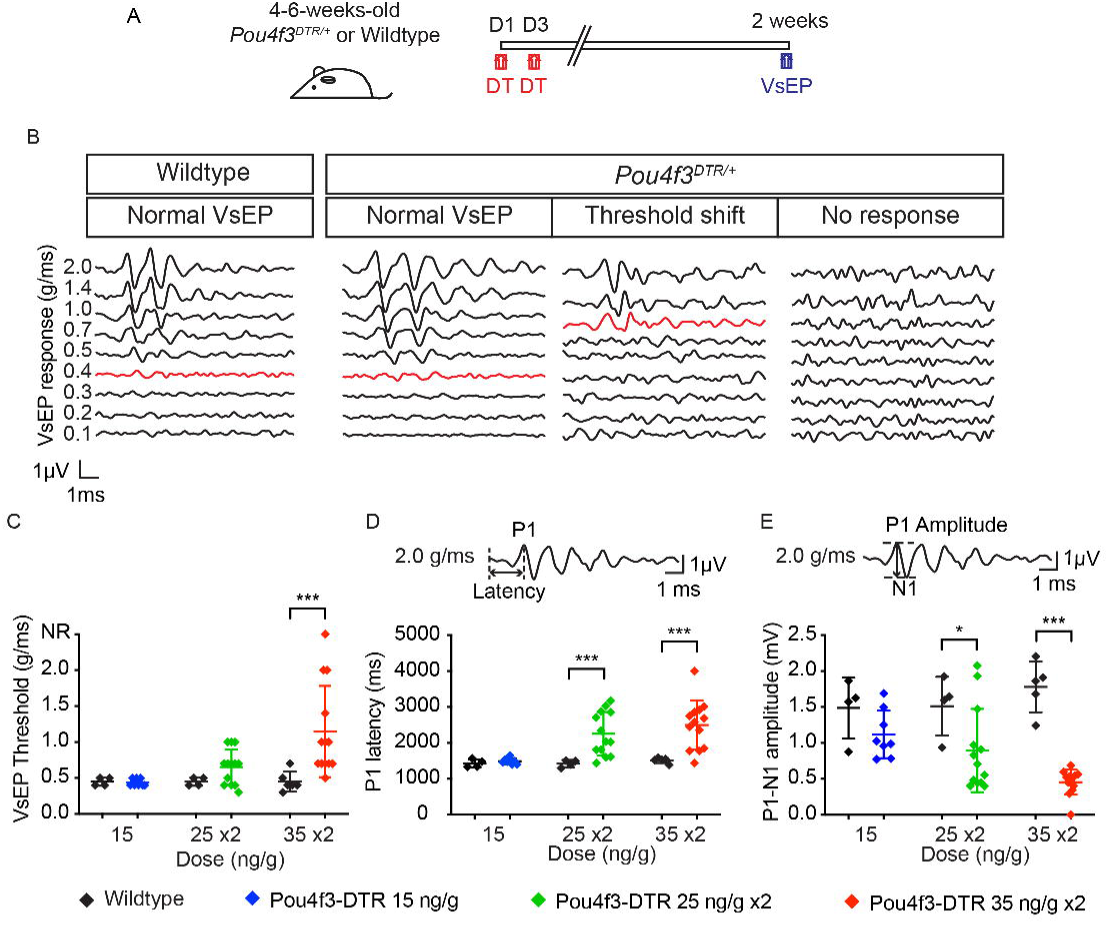
High dose diphtheria toxin treatment reduced VsEP responses of adult *Pou4f3^DTR/+^* transgenic mice. **A.** The experiment timeline. **B.** Representative VsEP waveforms are shown for a wild-type mouse (left), demonstrating a normal VsEP response with threshold marked in red, and for *Pou4f3^DTR/+^* mice (right), which may exhibit a normal response, an elevated threshold, or a complete absence of VsEP response (thresholds marked in red). **C-E.** Quantification of VsEP response threshold (B), P1 latency (C), and P1–N1 amplitude (D) in wild-type and FVB *Pou4f3^DTR/+^* mice treated with varying doses of diphtheria toxin. Significant shifts in threshold, increased P1 latency, and reduced P1–N1 amplitude were observed in *Pou4f3^DTR/+^* mice following high-dose DT treatment (35 x2 ng/g), but not with low doses (15 or 25 x2 ng/g). Data shown as mean±SD, and were analyzed using two-tailed, unpaired Student *t* tests. ***p<0.001, **p<0.01, *p<0.05. n=4 mice for wild type, and n=8 13, and 13 mice for *Pou4f3^DTR/+^*, treated with 15, 25 x2, and 35 x2 ng/g DT, respectively.

The effects of DT administration on otolith function were further assessed by measuring the VOR responses to sinusoidal head translation (tVOR) after 2 DT regimens (25 ng/g x2 and 50 ng/g x2). These regimens led to 27% and 19% hair cell survival in the extrastriola, and 24% and 17% hair cell survival in the striola (Table 2). At 2 weeks post DT injection, tVOR gains, sensitivity and phase were largely intact compared to undamaged controls at either dose (Figure 6A-D’).

**Figure 6.**
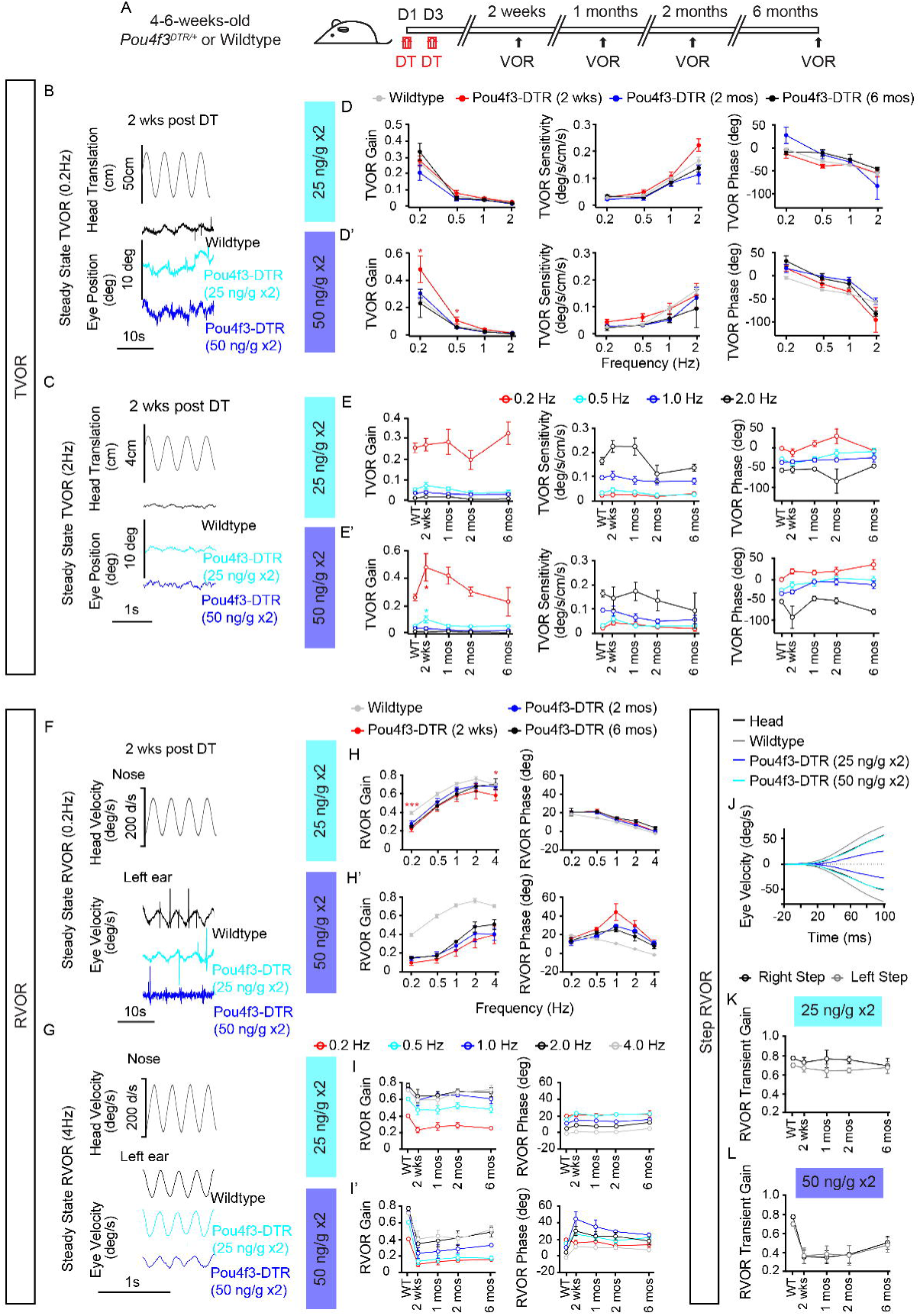
Effects of DT injections on the translational VOR (tVOR) and rotational VOR (rVOR). **A.** The experimental timeline. **B-C.** Representative eye movement responses to head translation at 0.2Hz (B) or 2Hz (C) in wild-type mice and 4-6-week-old C57BL/6J *Pou4f3^DTR/+^* at two weeks post DT treatment of 25 ng/g x2 or 50 ng/g x2**. D-D’.** Effects of DT treatment of 25 ng/g x2 (D) or 50 ng/g x2 (D’) on tVOR gains (left panels), sensitivities (middle panels), and phases (right panels) as functions of head translation frequency at two weeks, 2 months, and 6 months post DT treatment. **E- E’.** tVOR gains (left panels), sensitivities (middle panel), and phases (right panels) as functions of the interval post DT injection of 25 ng/g x2 and 50 ng/g x2, respectively. **F-G.** Representative eye movement responses to head rotation at 0.2Hz (F) or 4Hz (G) in a control mouse and mice at two weeks post DT treatment of 25 ng/g x2 or 50 ng/g x2. **H-H’.** Effects of DT treatment of 25 ng/g x2 (H) or 50 ng/g x2 (H’) on steady state rVOR gains (left panels) and phases (right panels) as functions of head rotation frequency at two weeks, 2 months, and 6 months post DT treatment. **I-I’.** Steady state rVOR gains (left panels) and phases (right panels) as functions of the interval post DT injection of 25 ng/g x2 and 50 ng/g x2, respectively. **J.** Averaged step rVOR responses at two weeks post DT treatment of 25 ng/g x2 (light blue) or 50 ng/g x2 (dark blue). **K-L.** Step rVOR gains as functions of the interval post DT treatment of 25 ng/g x2 and 50 ng/g x2, respectively. Data shown as mean±SEM, and were analyzed using two-tailed, unpaired Student *t* tests. ***p<0.001, **p<0.01, *p<0.05. n=19 mice for wild type, n=5, 5, 4, and 3, 4, 4, 4, and 2 mice for *Pou4f3^DTR/+^* at 2 weeks, 1 month, 2 months, and 6 months, treated with 25 x2 and 50 x2 ng/g DT, respectively.

The effects of DT administration on the crista/canal function were assessed by measuring the rVOR responses in two ways: first to sinusoidal head rotation (steady state rVOR) (Figure 6A, F-H’) and second, to step head rotation (step rVOR) (Figure 6J-L). While DT at 25 ng/g x2 resulted in hair cell survival of 31.0% in both the peripheral and central regions of the cristae, the steady state rVOR exhibited a reduction of 45.0% in gain at 0.2 Hz (p<0.001) and only 14.3% at 4Hz (p<0.05), with no significant change in phase at both frequencies (Figure 6F-H). This suggests that the low frequency response was more severely affected after hair cell loss. At higher DT doses (50 ng/g x2) where we detected hair cell survival at 18% and 5% in the peripheral and central regions, respectively, rVOR gain decreased 75.0% (p<0.001) and 42.8% (p<0.001) at 0.2 and 4 Hz, respectively, again with little phase changes at both frequencies but the low frequency was more severely affected (a phase lead was observed at 1 Hz) (Figure 6F-G, H’). Similar results were observed in step rVORs (Figure 6J-L). These results suggest a dose-dependent relationship between hair cell integrity and crista/canal function. Similar to the utricle, a substantial loss of hair cells only caused a small and partial loss of organ function.

In both the utricle and anterior canal, we have observed spontaneous regeneration of hair cells mainly in the extrastriolar/peripheral regions (Fig. 3G-H, 4G-H). We measured tVORs and rVORs at 1, 2, and 6 months post DT treatment (Figures 6E-E’, I-I’), while we found no significant improvement at either of the two DT doses, an increasing trend was observed over time (2 weeks, 1 month, 2 months, and 6 months).

### DT-induced hair cell loss causes deficits in vestibular afferents

We next performed single-unit recording from vestibular afferents, which is deemed a sensitive assay for vestibular function beyond reflex-based assays, during spontaneous firing and in response to head rotation and translation in DT-treated animals. We measured a total of 198 vestibular afferents in 5 DT-treated mice (3 months post DT treatment at 25 ng/g x2) and 195 afferents in 4 undamaged, control mice (Figure 7A, Table 4). Vestibular afferents are classified as regular or irregular coefficient of variation (CV*<=0.1 or >0.1, respectively) (Goldberg 2000, Lasker, Han et al. 2008) (Figure 7C, E and G). DT significantly reduced spontaneous firing rates of both regular (control: 73.9±1.9 spike/s, n=106; DT: 51.7±3.1 spike/s, n=56; p<0.01) and irregular afferents (control: 60.7±4.7spike/s, n=89; DT: 49.2±3.2 spike/s, n=142; p<0.05) (Figure 7B-B’, D-D’, F-F’, Table 4), consistent with damage to hair cells in both the extrastriolar/peripheral and striolar/central regions of both the cristae and macular organs, respectively. Interestingly, DT treatment significantly decreased the regularity of spontaneous firing of the regular afferents (control: 0.0503±0.0017, n=106; DT: 0.0665±0.0025, n=56; p<0.001) while having little effect on the regularity of irregular afferents (Figure 7C-C’, E-E’, G-G’, Table 4). We hypothesize that this decrease in CV* may cause some pre-DT injection regular afferents to be re-classified as irregular afferents. In support of this notion, DT treatment significantly reduced the proportion of regular afferents (control: 54.5% (106/195) versus DT: 28.3% (56/198), p<0.001) (Table 4). Moreover, DT treatment significantly reduced sensitivity to head rotation in the regular canal afferents (p=0.043), but not in the irregular canal afferents (Figure 7L-M). DT treatment also increased the distortions of the regular canal afferents to head rotation (P=0.001) (Figure 7L, O, Table 4). As expected, DT treatment did not reduce otolith regular afferents’ sensitivity to head translation in the naso-occipital direction. Instead, a trend toward an increase in sensitivity was observed (Figure 7H-K, Table 4). Nevertheless, we observed an increasing trend in otolith afferents’ distortion to head translation (Figure 7H, K). Furthermore, we found a significant increase in the proportion of unresponsive afferents in the DT-treatment group (DT-treated mice: 39%, 78/198 versus control mice: 17%, 34/195, chi-square=22.81, p<0.001, Table 4). In summary, DT treatment decreased spontaneous discharge rates of vestibular afferents, their responsiveness to head rotation, but no significant effects on their responsiveness to head translation.

**Figure 7.**
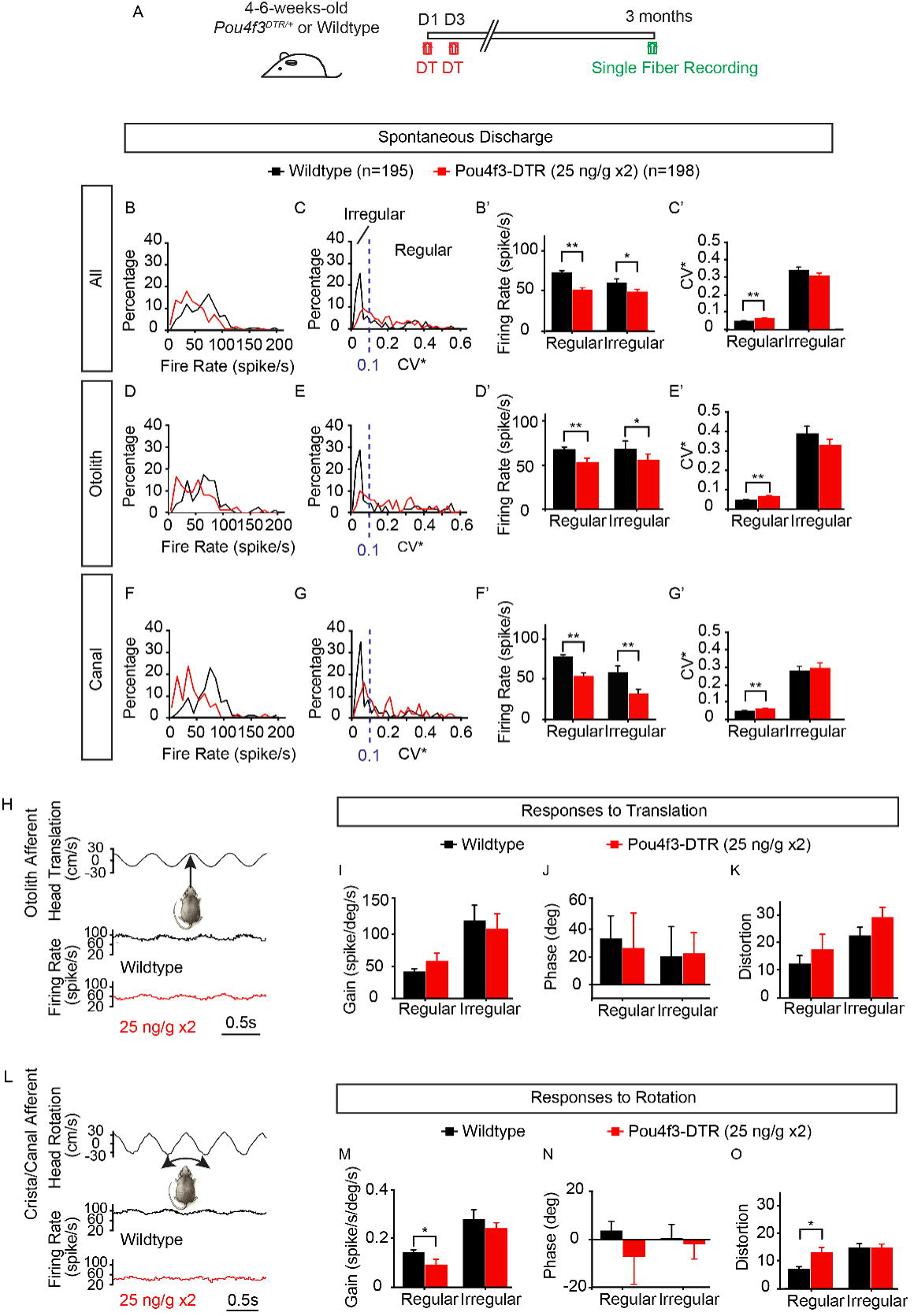
Otolith and canal afferent function. **A.** The experimental timeline. **B-G’.** Effects of DT treatment on all afferent spontaneous firing rates and regularity (B-C’), otolith and canal afferent spontaneous firing rates and regularity (D-G’). **H.** Representative responses to head translation of otolith afferents from a control mouse and a mouse at 3 months post DT treatment of 25 ng/g x2. **I-K.** Effects of DT treatment on otolith afferents’ sensitivities, phases, and distortions to sinusoidal head translation (1Hz) in the nasal-occipital direction, respectively. **L.** Representative responses to head rotation of canal afferents from a control mouse and a mouse at 3 months post DT treatment of 25 ng/g x2. **M-O.** Effects of DT treatment on canal afferents’ gains, phases, and distortions to sinusoidal head rotation (1Hz), respectively. Data shown as mean±SEM, and were analyzed using two-tailed, unpaired Student *t* tests. **p<0.01, *p<0.05. n=195 afferent recordings from 4 wild type and 198 from 5 *Pou4f3^DTR/+^* mice.

## Discussion

### Hair cell loss and vestibular dysfunction across different damage models

Several ototoxins have been used to induce hair cell loss and vestibular dysfunction in experimental animals. For example, aminoglycosides cause time- and dose-dependent loss of vestibular hair cells, particularly targeting type I hair cells, resulting in significant functional deficits (Lyford-Pike, Vogelheim et al. 2007, Sultemeier and Hoffman 2017). Cisplatin also induces vestibular hair cell loss, with the greatest damage observed at moderate doses, while higher doses paradoxically result in reduced toxicity (Ding, Jiang et al. 2018). Though not used clinically, IDPN also causes a dose-dependent loss of type I hair cells, with lesser effects on type II hair cells even at high doses, resulting in severe vestibular deficits (Llorens, Dememes et al. 1993, Sayyid, Wang et al. 2019, Zeng, Ni et al. 2020, Tian, Huang et al. 2024, Wang, Yang et al. 2024, Louise Schenberg 2025). However, these drugs (particularly IDPN) have been reported to cause collateral damage to the peripheral and central nervous systems in addition to the inner ear sensory cells, thereby further contributing to functional impairment.

The Pou4f3-DTR mouse is a widely used genetic model for selectively ablating hair cells in the auditory and vestibular organs (Cox, Chai et al. 2014, Atkinson, Dong et al. 2018, Kim, Wang et al. 2022). Previous studies using this transgenic model have shown that loss of over 90% of vestibular hair cells leads to significant vestibular dysfunction (Jauregui, Scheinman et al. 2024, Lahlou, Zhu et al. 2024). In our study, we demonstrated a dose-dependent loss of both type I and type II hair cells in both the anterior crista and utricle, even when overall hair cell loss reached more than 80%. Relative to other ototoxins, DT induces less damage to type I hair cells. Using VsEP, tVOR, as well as single unit recording to assess otolithic function, we found that utricle function is largely preserved until hair cell loss exceeded 90%. By contrast, measurements using rVOR and single unit recording suggest loss of canal function is detected as early as ∼68% hair cell loss, with worsened function resulting from more severe loss of hair cells. Remarkably, with ∼90% hair cell loss in both the crista and utricle, responses remain detectable in most animals tested. These results are important as they delineate the redundancy of hair cells of the vestibular organs, with a greater level of redundancy observed for the macular organs relative to the crista ampullaris.

In the inner ear, the vestibular labyrinth contains five paired end organs—three semicircular canal cristae and two otolith maculae. Each end organ shows zonal specialization (central/peripheral in cristae; striolar/extrastriolar in maculae) and is innervated by two classes of vestibular afferents (regular and irregular) that contact two hair-cell subtypes (type I and type II) (Goldberg 2000, Eatock and Songer 2011). Because some behavioral and physiological assays target and are more sensitive to individual parts of this complex vestibular system, we sought to characterize these assays after an escalating dose of DT to induce an increasing degree of hair cell loss in the cristae and maculae. The results from steady state rVOR, step rVOR, steady state tVOR, VsEP, and single afferent recordings provided a comprehensive picture of DT-induced functional deficits. For example, while DT treatment at a low dose of 25 ng/g x2 had little effect on the steady state tVOR and high frequency rVOR, it resulted in a significant reduction in rVOR gain at 0.2 Hz, suggesting different vestibular organs have different degrees of redundancy that can be revealed, particularly at a lower frequency. Furthermore, single-unit recordings show that DT-induced hair cell loss resulted in significant reductions in spontaneous firing rates, sensitivities to head rotation, and increases in distortions to head rotation. Interestingly, DT-induced hair cell loss also caused regular afferents to fire less regularly than controls. Because spontaneous discharge regularity can be mediated by postsynaptic mechanisms (Goldberg 2000), these results suggest that DT not only reduces inputs from HCs to afferents but may also indirectly affect postsynaptic processes. Future computational modeling could yield a deeper understanding of the underlying mechanisms.

### Hair cell redundancy in the vestibular system

In zebrafish lateral-line neuromasts, while all hair cells are mechanosensitive, most remain synaptically silent under normal conditions. Upon damage, the synapses of these silent hair cells rapidly activate, restoring sensory function and serving as a reserve population to maintain sensory function (Zhang, Li et al. 2018). This suggests a potential mechanism for redundancy and repair in lower vertebrates. However, little is known about whether a similar reserve population exists in the mammalian inner ear. Recently, Schenberg and colleagues reported that some vestibular function is still detectable after ∼50% loss of hair cells caused by IDPN treatment, whereas losses above 80% abolish normal function. Notably, type I hair cell loss strongly correlated with VOR deficits (L, Simon et al. 2025). Similarly, our results indicate that tVOR responses remained preserved until around 80% of hair cells were ablated, while measurable deficits in VsEP and tVOR only emerged after hair cell loss exceeded approximately 90%. Together, these results suggest significant functional redundancy in the vestibular system, particularly the macular organs; physiological impairments are only detected when a critical threshold of hair cell loss is surpassed. Future work should examine whether a silent or reserve population of hair cells exists in the mammalian vestibular system and explore their potential as a therapeutic strategy for restoring vestibular function.

### Divergent innervation of utricular hair cells by single afferents

In the vestibular system, afferent neurons from the vestibular ganglion transmit head movement information to the brain by innervating multiple hair cells through diverse patterns. Unlike cochlear inner hair cells, vestibular afferents commonly branch and contact several hair cells, enabling broad integration of sensory input. These afferents are categorized as calyx, bouton, or dimorphic fibers. Calyx afferents form large terminals around 1 to 3 neighboring type I hair cells in striola/central zones, bouton afferents make 15 to 100 contacts with type II hair cells in extrastriola/peripheral zones, and dimorphic afferents, the most common, exhibit mixed innervation with 1 to 4 calyces on type I hair cells and 1 to 50 boutons on type II hair cells across sensory epithelium (Goldberg 2000). This divergent innervation pattern allows individual afferents to gather input from multiple hair cells across spatial zones, allowing for robust signal transmission. However, whether each hair cell contributes equally to the afferents is not known. Thus, it is conceivable that when a significant number of hair cells are lost, these afferents may still receive input from surviving hair cells, which help preserve afferent responses to head rotation or translation. Such an anatomical arrangement may contribute to functional redundancy in the vestibular system, allowing it to maintain function even after substantial hair cell loss.

### Vestibular function is evolutionarily vital for survival

The vestibular system is evolutionarily ancient and essential for survival, with its basic structure established early in vertebrate evolution, even in jawless species. Unlike the auditory organ, which appeared later and has undergone significant evolution, the vestibular system developed as a fundamental adaptation to support balance, gaze stabilization, and spatial orientation. The five vestibular organs are highly conserved from fish to mammals, highlighting their essential role in coordinating eye and body movement in a three-dimensional environment (Straka, Zwergal et al. 2016, Mackowetzky, Yoon et al. 2021). In many teleost fish, the vestibular epithelia also serve auditory functions due to the absence of a separate auditory organ. A distinct auditory structure first appeared in amphibians and later evolved into the highly specialized cochlea in mammals. These evolutionary trajectories suggest that effective vestibular function was achieved early and preserved due to its essential role in survival. Given the fundamental importance of vestibular function to the organism, vestibular function may therefore have developed fail-safe mechanisms to preserve its function in the face of mild insults.

In summary, we have shown through the use of the Pou4f3-DTR mouse model that DT treatment can induce a dose-dependent loss of hair cells in the crista ampullaris and utricle. Using a battery of vestibular-specific tests (tVOR, rVOR, VsEP, and single unit recording), we showed that utricle function is largely preserved until hair cell loss exceeds ∼90%, whereas abnormal crista function is detected when loss exceeds ∼70%. These findings should inform future work on vestibular repair/regeneration and possibly tests on human vestibular dysfunction.

## Methods

### Mice

*Pou4f3^DTR/+^* transgenic mice (The Jackson Laboratory, #028673, Bar Harbor, Maine, USA), wild type C57BL/6J background (The Jackson Laboratory, #000664), and wild-type FVB/NJ (The Jackson Laboratory, #001800) were used. All experimental Pou4f3-DTR C57BL/6J mice were maintained as a pure inbred line. FVB-background Pou4f3-DTR mice were generated by backcrossing C57BL/6J *Pou4f3^DTR/+^* mice with wild-type FVB/NJ mice for over ten generations. Mice of both sexes were used. Animals were housed under standard husbandry conditions with open access to food and water. Both *Pou4f3^DTR/+^* and wildtype littermates (4-6 weeks old) were injected with diphtheria toxin (DT; Millipore Sigma, #322326, Burlington, Massachusetts, USA, and List Lab, #150, Campbell, California). DT doses (intramuscular, IM) ranged from a single dose of 15 ng/g, 25 ng/g x2, 35 ng/g x2, and 50ng/g x2, with double doses spaced 2 days apart. During the first week after the DT injection, some FVB mice received 0.5 ml of lactated Ringer’s solution (subcutaneous 2-3 times daily and a high-calorie gel was supplemented (Suppli-Cal, Henry Schein, #029908). The Animal Care and Use Committees of the Stanford University School of Medicine, the University of Mississippi, and NIH approved all protocols.

### Genotyping

Genomic DNA of transgenic mice was collected to perform PCRs. DNA was isolated by adding 180 μL of 50 mM sodium hydroxide (NaOH, Thermo Fisher Scientific, S318-500) to tissue biopsies, and incubating at 98°C for 1 hour, at which point 20 μL of 1 M Tris-HCl (Invitrogen, Thermo Fisher Scientific, #15568025) was added. The following primers and sequences were used for genotyping. Mutant Forward: 5’-GTCAAAAAATGTGCCTTAGAG -3’; mutant reverse: 5’-CCGACGGCAGCAGCTTCATGGTC-3’; wildtype forward: 5’-CACTTGGAGCGCGGAGAGCTAG-3’.

### Vestibular physiology

Linear vestibular evoked potential (VsEP) responses were recorded from mice as previously described (Jones and Jones 1999). Briefly, mice were first anesthetized with a 1:1 cocktail of ketamine (100 mg/kg, #NDC 50989-161-06, Vedco, St. Joseph, Missouri, USA) and xylazine (10 mg/kg, #SC-36294Rx, Santa Cruz Animal Health, Dallas, TX, USA). Rectal temperatures were maintained at 37°C, and electrocardiographic (ECG) activity was monitored using an oscilloscope. Subcutaneous stainless-steel electrodes were placed over the caudal cerebrum (non-inverting electrode), subcutaneously behind the right pinna (inverting electrode), and intramuscularly in the right thigh muscle (ground electrode). A head clip was used to secure the head to the mechanical shaker, which was used to deliver linear vestibular stimuli in the naso-occipital axis. Vertical motion of the shaker was monitored with an accelerometer and adjusted to produce the stimulus waveforms. Throughout the study, jerk stimuli ranging from 0.125 to 2.0 dB were provided (Jones, Jones et al. 2011). Signals were amplified (200,000x), filtered (low filter = 300 Hz, high filter = 3000 Hz), and digitized via an analog-to-digital (A/D) converter for all VsEP recordings. Responses from normal and inverted stimulus polarities were collected and added together for a total of 256 sweeps for each waveform. A masker (90 dB SPL; bandwidth 50 Hz to 50 kHz) from a free-field speaker driver was used to prevent responses from the auditory components of cranial nerve VIII. All responses were blindly analyzed for three components: threshold (g/ms), P1 latency (ms) and P1-N1 amplitude (μV). Thresholds were defined as the average stimulus intensity between the lowest stimulus intensity at which a response was observed and the following stimulus intensity that fails to elicit a response. Latencies were measured relative to the onset of the stimulus for the first positive response peak (P1). Amplitudes represented peak-to-peak magnitudes between P1 and N1. Whereas the first positive and negative response peaks (P1 and N1, respectively) reflect activity of the peripheral vestibular nerve, peaks beyond N1 reflect activity of the brainstem and central vestibular relays (Jones 1992, Nazareth and Jones 1998).

### Immunohistochemistry

Following harvesting, inner ear vestibular organs including utricle and anterior crista ampullaris were fixed for 40 minutes in 4% paraformaldehyde (in PBS, pH 7.4; Electron Microscopy Services, #15710) at room temperature. Tissues were rinsed with PBS 3X for at least 15 minutes each, then blocked with 5% donkey serum, 0.1% TritonX-100, 1% bovine serum albumin (BSA, Thermo Fisher Scientific, #BP1600-100), and 0.02% sodium azide (NaN_3_, Sigma-Aldrich, #S2002-25G) in PBS at pH 7.4 for 1 hour at room temperature. Primary antibodies diluted in the same blocking solution were added overnight at 4°C. The next day, after washing with PBS, tissues were incubated with secondary antibodies diluted in 0.1% TritonX-100, 0.1% BSA, and 0.02% NaN_3_ solution in PBS at pH 7.4 for 2 hours at room temperature. After washing with PBS, tissues were mounted in antifade Fluorescence Mounting Medium (DAKO, Agilent, #S3023, Santa Clara, California, USA) and coverslipped.

Antibodies against the following proteins were used: Myosin7a (1:1000, mouse, DSHB, RRID: AB_2282417), Myosin7a (1:1000, rabbit, RRID: AB_2314839, Proteus Biosciences), Sox2 (1:400, goat, RRID: AB_2286684, Santa Cruz Biotechnology). Secondary antibodies included Alexa Fluor donkey anti-goat 647 (RRID: AB_ 2535864, Thermo Fisher Scientific), Alexa Fluor donkey anti-rabbit 488 (RRID: AB_2535792, Thermo Fisher Scientific), Alexa Fluor donkey anti-rabbit 546 (RRID: AB_2534016, Thermo Fisher Scientific), at 1:250 to 1:500. Fluorescence-conjugated DAPI (1:10,000 from 5 mg/ml stock solution; RRID:AB_2629482, Thermo Fisher Scientific) were used.

### Image acquisition and cell quantification

Images were acquired using epifluorescent or confocal microscopy (LSM700 or LSM880, Zeiss) and analyzed with ImageJ (64 bit) and Fiji (NIH) (Schindelin, Arganda-Carreras et al. 2012) and Photoshop CS6 (Adobe Systems). Optical slices of 1 µm thick were used in z stack images. Low-magnification images were captured using 10x objective on epifluorescent microscope (Axioplan 2, Zeiss). High-magnification Z-stack images were acquired using a 63x objective on a Zeiss LSM 700 or 880 confocal microscope (Zen Black, Zeiss). From these Z-stack images, cell quantification was performed either per 10,000 μm^2^ (utricle), 2500 μm^2^ (crista ampullaris) or per organ using Fiji software (v2.0.0, NIH). Cell counts were analyzed either from 1-2 representative areas from the extrastriola and striola (utricle), peripheral zone and central zones (anterior crista ampullaris), or from merged images spanning the whole utricle (merged together using Zeiss Zen software). For all experiments, n values represent the number of animals or cells examined.

### Vestibulo-ocular reflexes (VORs)

Each mouse was implanted with a head holder on the skull as described before (Stewart, Yu et al. 2016, Lentz, Pan et al. 2020, Yu, Huang et al. 2020, Lahlou, Zhu et al. 2024, Du, Huang et al. 2025). Eye position signals of the left eye were recorded using an ISCAN eye tracking system mounted on a rotator/sled. The rotational VOR (rVOR) was tested at 0.2-4 Hz (60 deg/s peak velocity) and the translational VOR (tVOR) was tested at 0.2-2Hz (0.1g peak acceleration). To measure the transient rVOR responses, rapid leftward or rightward 10° head rotation steps were delivered reaching a velocity of 100°/s within 100 ms with a peak acceleration of ∼1500°/s/s. At least 30 cycles/50 trials per condition were recorded. Signals related to eye position and head position were sampled at 1 kHz by a CED system. Gains and phases of the VORs were calculated by performing fast Fourier transform (FFT) analysis.

### Single unit recording of vestibular afferents

Single unit recording of vestibular afferents was performed under ketamine/xylazine anesthesia as described before (Zhu, Tang et al. 2011, Lentz, Pan et al. 2020, Huang, Tang et al. 2022, Lahlou, Zhu et al. 2024). Briefly, a craniotomy was performed to allow access of the vestibular nerve by a microelectrode. Extracellular voltage signals were sampled by a CED at 20 kHz with 16-bit resolution and a temporal resolution of 0.01 ms. Head position signals were sampled at 1 kHz. Each afferent’s spontaneous activity was recorded for computing resting firing rates. Regularity of vestibular afferents was determined by calculating their normalized coefficient of variation (CV*) of inter-spike intervals. Vestibular afferents are classified as regular (CV*<=0.1) or irregular (CV*>0.1) units based on their CV* (Goldberg 2000, Lasker, Han et al. 2008). To quantify an afferent’s responses to head rotation and translation, the fundamental response was extracted from the averaged data using a fast Fourier transform (FFT) analysis. Gains and phases relative to head rotation velocity and translation acceleration were calculated at ∼1 Hz. Given that many afferents in damaged mice show minimal modulation during head movement, traditional gain metrics alone are insufficient for assessing signal significance in their discharge activities. To address this, we employed a distortion metric, defined as 1 - (amplitude of the fundamental response)/(square root of the sum of the squared amplitudes of the first 10 harmonics), for a more rigorous evaluation of head movement signals in vestibular afferents. This distortion metric, akin to “stimulus-response coherence”, is particularly effective for assessing vestibular afferent responses to head rotation and translation in animal models with damaged vestibular end organs resulting from genetic mutations, trauma and ototoxicity (Sultemeier and Hoffman 2017, Huang, Tang et al. 2022, Du, Huang et al. 2025). An afferent with translation distortion <=30% is classified as a translation/otolith afferent. An afferent with rotation distortion <=30% and translation distortion >30% is classified as a rotation/canal afferent. An afferent with rotation distortion >30% and translation distortion >30% is classified as a “no-response” afferent.

### Statistical analyses

Statistical analyses were conducted using GraphPad Prism (v7.0c software, GraphPad). Statistical significance was determined using unpaired Student *t* tests, or one- or two-way ANOVA with ordinary multiple comparison test unless otherwise stated as appropriate. Chi-square test was used to compare the proportion of ‘no-response’ and regular/irregular afferents. A p value less than 0.05 was considered significant: *p<0.05, **p<0.01, ***p<0.001. Data are shown as mean±SD or mean±SEM. Details of experiments can be found in the figures and figure legends. Numbers of animals or HCs listed in parentheses.

## Acknowledgements

We thank the Cheng laboratory for insightful comments on the manuscript and S. Jones for excellent advice. This research was supported by Stanford Medical Scholars Research Program, Howard Hughes Medical Institute Medical Fellows Program, Stanford MSTP, and NIH/NIDCD F30DC015698 (Z.N.S), Lucile Packard Foundation for Children’s Health, Stanford NIH-NCATS-CTSA UL1 TR001085, Child Health Research Institute of Stanford University, and NIH/NIDCD R21DC015879, National Natural Science Foundation of China 82071056 (T.W.), NIH/NIDCD R01DC018919 (H.Z.), 1P20GM156708-01 (W.Z.), K24DC020986, RO1DC020879, RO1DC021110 and RO1DC016919 (A.G.C.), and the Stanford Initiative to Cure Hearing Loss.

## Author contributions

T.W., D.K.H., Z.N.S., H.Z., W.Z., A.G.C. designed experiments, T.W., D.K.H., A.M., Z.N.S., J.H., C.S., Z.G.I., H.Z., W.Z., A.G.C. performed experiments, T.W., D.K.H., A.M., Z.N.S., J.H., H.Z., W.Z., A.G.C. analyzed data, T.W., D.K.H., A.M., Z.N.S., H.Z., W.Z., A.G.C. prepared the manuscript.

**Supplementary Figure 1. Characterization of the utricle sensory epithelium area and supporting cell density in the adult *Pou4f3^DTR/+^* mice**.

**A.** DT treatment causes a significant reduction of Myosin7a^+^ sensory epithelium area in utricle from *Pou4f3^DTR/+^* mice, particularly at high doses. **B-C.** Quantification of Sox2^+^ supporting cells shows no significant change in the extrastriolar (**B**) and striolar (**C**) regions following DT treatment at specified doses. Data shown as mean±SD, and were analyzed using two-tailed, unpaired Student *t* tests. ***p<0.001, **p<0.01, *p<0.05. n=8, 7, 4, and 6 mice for wild type, and n=14, 18, 13, and 10 mice for *Pou4f3^DTR/+^*, treated with 15, 25 x2, 35 x2, and 50 x2 ng/g DT, respectively.

## Notes

### Competing Interest Statement

The authors have declared no competing interest.

### Summary of Updates

Paper is being resubmitted to a journal

