## Supplementary figures and images for "Graded Hair Cell Ablation Reveals Functional Redundancy in the Mature Mouse Vestibular System"

### Supplementary figure 1.tif

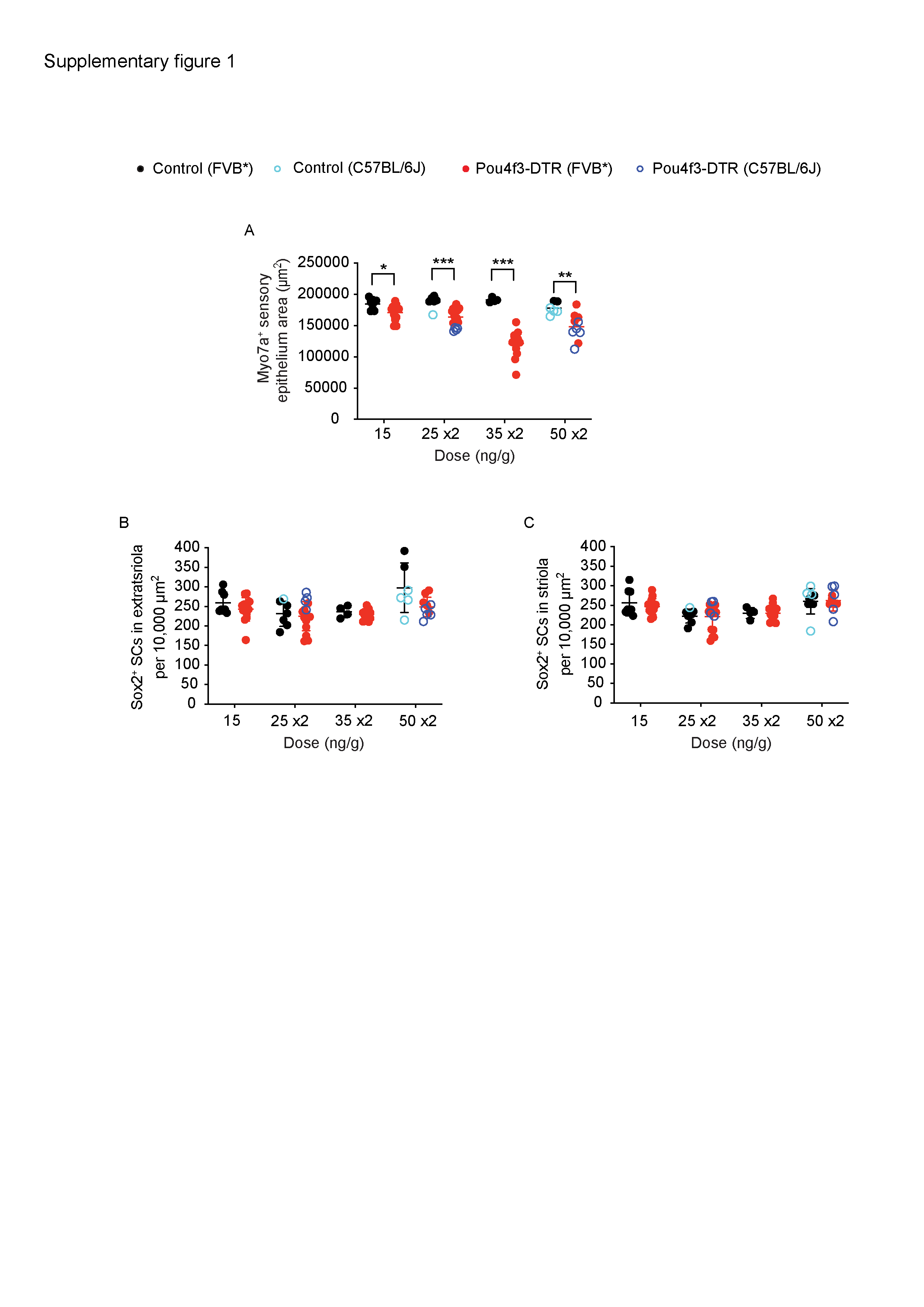
